# Evolutionary history of ligand binding by the LRR domain of innate immunity receptors: the story of the TLR2 cavity

**DOI:** 10.64898/2026.03.26.714386

**Authors:** Rami Namou, Keita Ichii, Abraham Takkouche, Lukasz Jaroszewski, Adam Godzik

## Abstract

Toll-like receptors (TLRs) are vital components of the innate immune system, recognizing both exogenous pathogens signals (PAMPs) and internal stress signals (DAMPs). TLR2 is unique among the human (*Homo sapiens*) TLR family members, as it contains a large cavity for binding hydrophobic ligands, such as lipoteichoic acid (LTA) and di/triacyl lipopeptides (Pam2/3CSK4). This study analyzed the structural phylogeny of cavity presence in the TLR2 lineage in vertebrates (vTLR) enabled by AI protein structure predictions and explored the potential convergent evolution of similar features in invertebrates (iTLRs).

Analysis of AI models of TLR2s shows that this cavity is consistently present in TRL2 orthologs across jawed vertebrates (Gnathostomata). In jawless vertebrates (Cyclostomatha), these cavities were found in lamprey (*Petromyzon marinus*) TLR2 model, but only in some extant hagfish (*Myxini*), suggesting an ancestral origin in basal vertebrates followed by lineage-specific losses. TLR2 paralogs were found in several species, with a similar central cavity but potentially different ligand specificities. In silico ligand docking showed Pam2CSK4 binds to this cavity in all TLRs and paralogs consistently, demonstrating the conserved function of the ligand-binding pocket in gram-positive bacteria recognition across TLR2 branches. Changes in the TLR2 cavity size and shape in some vertebrate groups show the evolution of this DAMP recognition mechanism adapted to its respective pathogens.

iTLRs form a separate phylogenetic branch with distinct structural features, but in literature some are considered to be TLR2 orthologs. Indeed, TLRs from some species of *Helobdella* and *Ciona*, contain a cavity with some similarity to that in the vTLR2 lineage. However, detailed structural comparisons of their location in the LRR domain and the structural details of the models suggest that their cavities have developed independently from that in TLR2s. Smaller cavities are present in other branches of the LRR family, but show different locations, shapes, and features, indicating that the binding of small ligands in the internal cavities within the LRR domains evolved multiple times in the LRR domain family history.

## Introduction

Toll-like receptor 2 (TLR2) plays a distinctive role within the Toll-like receptor (TLR) family as the primary sentinel of the innate immune system response to gram-positive bacterial pathogens, which are responsible for a significant proportion of the global infectious disease burden. It also recognizes endogenous ligands released during tissue stress or injury. Various components of the innate immune system, including neutrophils, macrophages, dendritic cells, eosinophils, and natural killer cells, express TLR2 [1].

The structures of many vTLRs and iTLRs have been determined at atomic resolution and deposited in the Protein Data Bank (PDB), including those of mouse (*Mus musculus*) and human TLR2 [2,3]. They all contain a leucine rich repeat (LRR) domain with a characteristic curved solenoid shape, acting as a binding domain recognizing specific ligands. Experimental studies have shown that TRL2s cavities serve as the binding site for several key TLR2 agonists representing different pathogens as well as endogenous DAMPs, including lipoteichoic acid (LTA) (Gram-positive bacteria), peptidoglycan (Bacteria), zymosan (Yeast/Fungi), Pam2CSK4 (Gram-positive bacteria), and Pam3CSK4 (Gram-positive bacteria). Additional experimentally validated TLR2 agonists include chitin (fungi), heat shock protein 60 (Hsp60), glycolipids (bacteria), glycosylphosphatidylinositol (GPIs) (bacteria), porins (bacteria), arabinogalactan (mycobacteria), lipomannan (mycobacteria), and lipoproteins such as LprA and LpqH (mycobacteria) [4], but their binding modes remain unknown. These ligands initiate robust TLR2-mediated inflammatory responses, underscoring the importance of ligand recognition in innate immunity.

Recent advances in AI-based protein modeling, such as AlphaFold2/3 [5,6], RoseTTAFold[7], and ESM2/3[8,9], have enabled accurate prediction of protein structures, providing an opportunity to use protein structure analysis tools on the complete repertoire of proteins, including TLRs (and TLR2s), across the genomes of multiple species. Structural features, such as cavities or unusual loops in structures that cannot be easily identified from amino acid sequences alone, are not included in classical phylogenetic analyses. This provides a unique opportunity to compare TLR2 cavities between humans and other vertebrate species to study their functions and evolutionary origins. Other TLRs contain much smaller cavities, such as TLR1’s cavity binding one long fatty acid (palmitoyl) chain [3] (versus two for TLR2) and TLR7/8’s two small pockets binding a guanosine/uridine and short RNA fragments [10,11]. Overall, the presence of a sizable cavity within an LRR domain is a relatively rare occurrence, but is still found in several other distant families of LRRs in animals (NLRs) [12] and plants (LRR receptors [13] and LRR F box proteins[14]).

Analysis of the ligand-binding cavities in the TLR2 lineage and other LRR proteins raises the broader question of how novel functional features emerge on conserved protein (scaf)folds. While overall folds and broadly defined functions are usually ancestral and are only slightly modified during the evolution of protein families, localized functional features such as metal-binding sites, active sites, and potentially cavities depend on a few specific positions in the sequence and could emerge independently in different branches of the same family by a process of convergent or parallel evolution. In our previous work, we identified such processes in the pathogenic adaptation of virulence factors [15] and carbohydrase kinase specificities [16], with many other similar examples reported in the literature [17,18]. To the best of our knowledge, such questions were never studied for ligand binding cavities in tandem repeat proteins.

## Methods

### Generation of vTLR, iTLR, and Human LRR protein datasets

The Human TLR2 gene corresponds to the Ensembl [19] entry ENSG00000137462 and Uniprot [20]: entry O60603. The gene tree of the TLR2 family was obtained from the Ensembl Gene Tree, and using Ensembl-UniProt equivalences, sequences and models for all proteins on this tree were downloaded from UniProt. If AF2 models were unavailable, AF3 models were calculated using the AlphaFold server. In particular, *Homo sapiens* (Humans) (O60603), *Mus musculus* (Mice) (Q9QUN7), *Gallus gallus* (Chicken) (Q9DGB6), *Anolis carolinensis* (G1KUN2), *Xenopus tropicalis* (A0A803JIX3), *Danio rerio* (Zebrafish) (F1QWS9) *Petromyzon marinus* (Pacific Sea lamprey) (S4S1Q8), and *Eptatretus atami* (brown hagfish) (ENSEBUG00000013629.1, A0A8C4QY81, model by Alphafold3). Ensembl-defined orthologs of the Human TLR2 gene were used as “vTLR” (vertebrate TLR dataset), 239 ortholog genes were identified, and modeled using Alphafold 3. To focus on the central and remove erroneous cavities elsewhere, another script extracted the LRR domain via removing the first 50 amino acids (signal domain) and last ∼150 amino acids (transmembrane segment and the TIR domain)

In the analysis of invertebrate TLR2, sequences were extracted from the proteomes of several protostomes. For *Ciona intestinalis*, the UniProt proteome UP000008144 was filtered using TIR and LRR keywords, yielding eight hits. For *Strongylocentrotus purpuratus*, the UniProt proteome UP000007110 was filtered using (Proteomecomponent:”Unassembled WGS sequence”) AND LRR AND TIR, resulting in 182 hits. For *Branchiostoma floridae*, the UniProt proteome UP000001554 was filtered using TIR and LRR, yielding 78 proteins. AF3 models were generated for both *Branchiostoma floridae* and *Saccoglossus kowalevskii* (acorn worm, 5 proteins), with the latter based directly on the FASTA files. In an evolutionary framework, the PANTHERDB database was queried using the term “TLR2.” Orthologs identified as “LDO” (Least Divergent Orthologs) were incorporated into the analysis. This approach identified the LeTLR2 (*Helobdella robusta)* (T1EUA2) and two duplicate *Ciona* genes Ci-TLR2 (*Ciona intestinalis*) (F6SGF2) and duplicate (C9K4U9). These five groups of invertebrate TLRs are known as “iTLR.”

Species-specific duplication events, present in only three lineages, were identified from the Ensembl TLR2 phylogenetic tree. The structural phylogeny of ancestral cavity features is depicted by comparing humans and lampreys to the convergently evolved cavity in the invertebrate leech. Duplicates of TLR2 were analyzed in chickens (C4PC57), green anoles (G1KWG2), and clawed frogs (A0A7D9NKA2).

Lastly, 325 proteins annotated as “LRR” in the human proteome according to Uniprot were identified using an in-house novel computational tool developed in our group that identified LRR domains from each full amino acid sequence, and the LRR domain sequences were extracted and modeled for each protein using Alphafold 3.

### PyVOL and Fpocket4 Calculation

The interactive versions of the PyVOL software were utilized to calculate volumes for all TLRs described (vTLRs, iTLRs and Human LRR proteins) This software was then used in batch mode and applied to AlphaFold3 models of TLR2 orthologues identified in the Ensembl database. Fpocket4 was used in an identical fashion to distinguish and compare two cavity calculation methods: one by fitting spheres (Fpocket 4) and the triangular mesh method (PyVOL). Fpocket4 was used for all vTLR and LRR human protein datasets.

### Ligand docking

Ligand docking illustrations were produced using Boltz 1 models accessed through a Neurosnap account[21,22]. Boltz 1 was employed to dock the ligands Pam2CSK4 and lipoteichoic acid onto TLR2 proteins modeled using AF3. The PDB outputs from Boltz 1 were subsequently input into the PRODIGY HADDOCK protein-ligand web server to compute the binding affinities [23]. LeTLR2 was uniquely docked with Pam2CSK4 to identify the binding cavities. Lipoteichoic acid (LTA) was also modeled using Boltz 1 and binding energy computed with PRODIGY, however due to dubious evidence of LTA physically docking to TLR2 is religated to Supp Info.

### Hagfish and Lamorey Ortholgue search

TLR2 orthologs from cyclostomes were identified using predicted protein sequences from ***Myxine glutinosa*** (Atlantic hagfish; XP_067991900, XP_067991085) and ***Eptatretus burgeri*** (inshore hagfish; A0A8C4QY81), together with additional lamprey taxa (NCBI Taxonomy IDs: 7757, 7755, 245074, 980415, 7748, 7750, 7753).

### MUSCLE Alignment

Atlantic hagfish (HgTLR2) (*Myxine glutinosa*), LeTLR2, CiTLR2 were aigned each with Human and Lamprey in three separate alignments using MUSCLE [24].

## Results

A comparison of the TLR2 cavities in all vertebrates showed a consistent shape and volume, as calculated by PyVOL [25] and confirmed by Fpocket4 [26,27] (Figure 1, above) (Table 1 in Supp. Info). Figure 1 (A–F) shows the central cavity in the LRR domain of the TLR2 protein in selected vertebrate species. The cavity in all vertebrate TLR2s is at the same relative location between positions 250 – 390 along the LRR domains with a volume of 1197 ± 46 A^3^. Its location coincides with the most divergent individual repeats, a sign of functional adaptation across the LRR domains, as discussed previously [28](preprint). The average TLR2 cavity volume across 10 manually analyzed vertebrate TLR2s was 1404 (±244.35) A^3^. The chicken (*Gallus gallus domesticus*) (1775.42 A^3^) and green anole (*Anolis carolinensis*) least divergent orthologs (LDOs) (1494.42 A^3^) were more than 1 SD away from the mean, indicating an expansion of the central cavity relative to that of other vertebrates. In contrast, the chicken TLR2 paralog (1029.21 A^3^) and lamprey TLR2 (1107.52 A^3^) were markedly smaller, by approximately 375 and 296 A^3^, respectively. The human (1334.25 A^3^) and mouse TLR2 (1177 A^3^, as referenced) are moderately below the average but remain within one standard deviation of the mean value. Such variation may be indicative of lineage-specific selective pressures due to different environmental niches, highlighting the evolutionary dynamics that shape the TLR2 ligand-binding cavity in vertebrates or an artefact of using AI based prediction algorithms. However, their shape and position within the LRR domain clearly identify them as descendants of the common ancestral TLR2.

**Figure 1:**
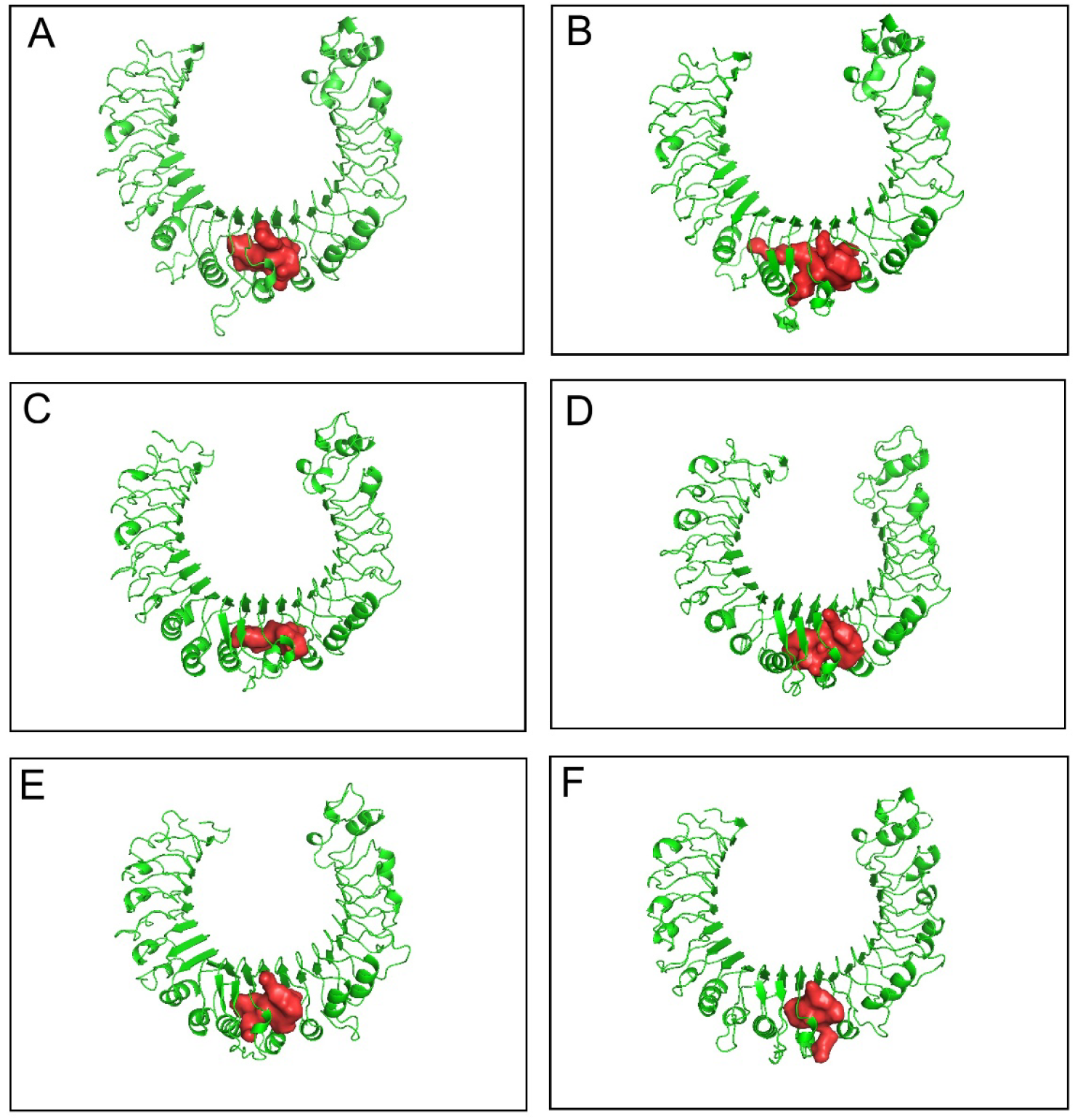
Comparison of the central cavity among 6 manually curated vertebrate’s TLR2s (A) *Homo sapiens*, (B) *Mus musculus*, (C) *Gallus gallus* (chicken) (LDO), (D) *Anolis carolinensis* (green anole)(LDO), (E) *Xenopus tropicalis* (western clawed frog)(LDO), (F) *Petromyzon marinus* (sea lamprey))

We hypothesize that the TLR2 ligand-binding cavity emerged either at the base of vertebrates or earlier in the lineages of invertebrates that branched off immediately preceding the emergence of vertebrates [29,30]. Figure 2 shows a structural comparison of the TRL2 cavities in humans, lampreys (*Petromyzon marinus*), and hagfish (*Eptatretus burgeri*), the last two representing jawless fish (Cyclostomatha, the only surviving clade of Agnatha), the basal phylum of vertebrates. The TLR2 LDO in the only hagfish represented in Ensembl and PantherDB does not have a large central cavity, in contrast to all other vertebrates. On the other hand, lamprey TLR2 LDO has a well-defined cavity; therefore, the most parsimonious scenario suggests that the common vertebrate ancestor possessed this feature, which was progressively lost in the hagfish lineage. This observation is further supported by the presence of much smaller cavities in the analogous TLR2 region in genomes of three other hagfish, which were not present in the PantherDB [31,32] or Ensembl [19] databases (see the Discussion section, for more analysis on lamprey and hagfish TLR2s)

**Figure 2A:**
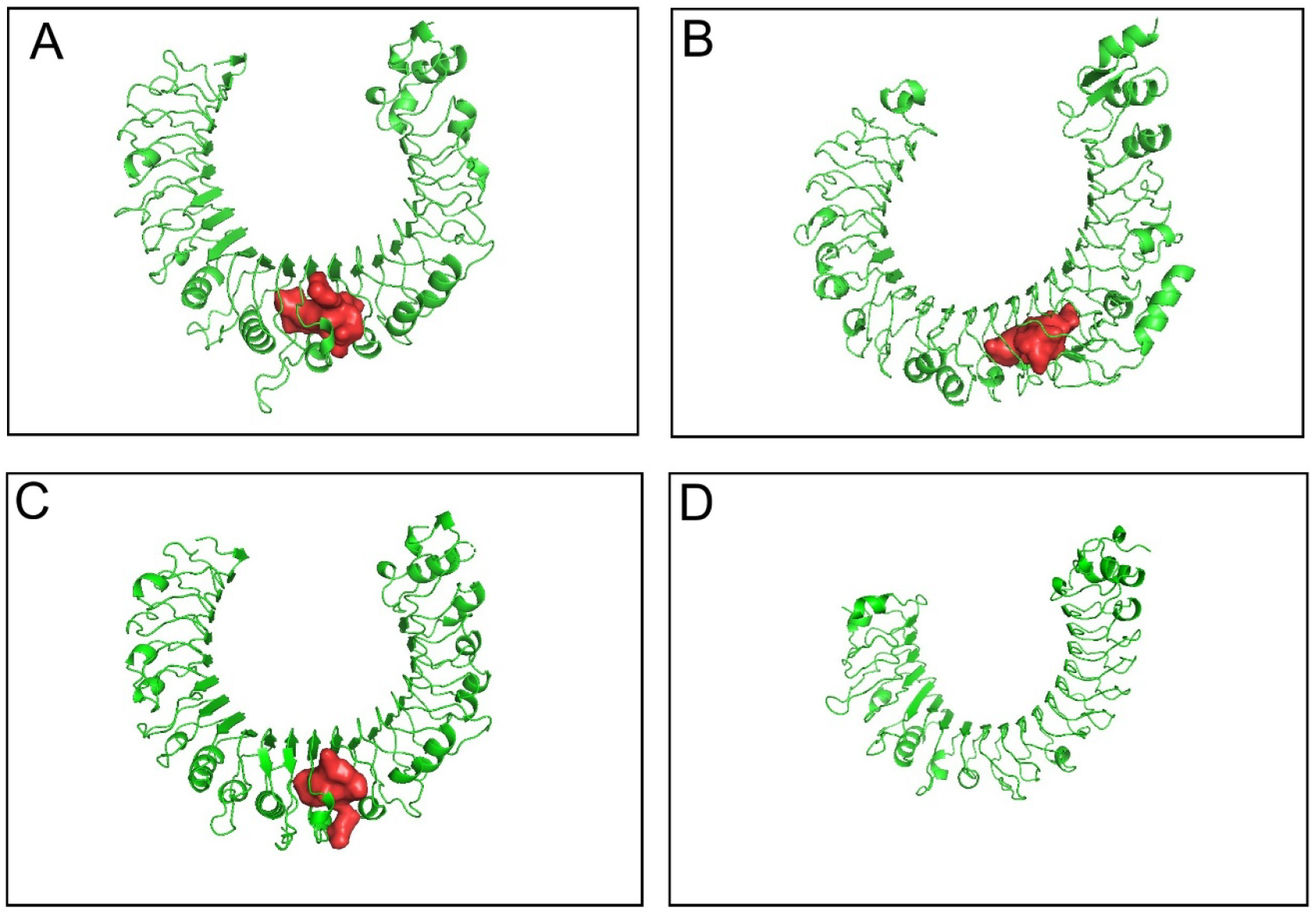
Structural comparison of TLR2 cavities in invertebrates and vertebrates. **(A)** *Homo sapiens* (human) **(B)** *Helobdella robusta* (california leech), **(C)** *Petromzon* marinus (sea lamprey), **(D)** *Eptatretus burgeni* (hagfish)

**Figure 2B:**
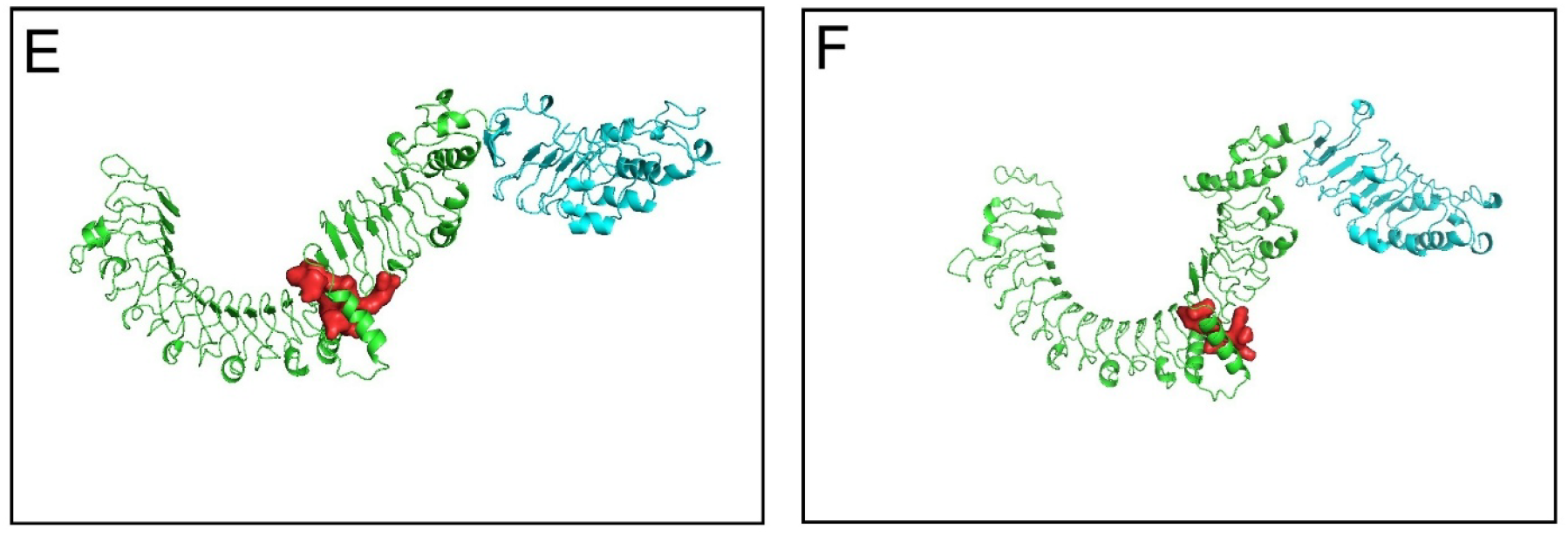
**B:** Comparison of the two *Ciona intestinalis* (vase tunicate) putative TLR2 paralogs **(E)** LDO and **(F)** duplicate.

Two invertebrate TLRs were annotated in the PantherDB and Ensembl as TLR2 orthologs and we analyzed them separately. Leech TL2 (LeTLR2) from *Helobdella robusta* was compared to vTLRS, humans, lampreys, and hagfish. LeTLR2 has a vTLR like structure with a single LRR domain with a cavity in the center and a TIR domain; thus, it could be the earliest example of a large central cavity in the TLR2 lineage. LeTLR2 can be aligned with human and lamprey TLR2s and, according to this alignment, the cavity region is located to the same regions between positions 300 – 450 along the LRR domain where and human and lamprey TLR2s cavity is located (S3C in Supp. Figures). Its cavity volume of 1195 A3 (generated with Boltz 1, Pam2CSK4 docked) was very similar to and within the average of vTLR (1197 ± 45.66 A3). However, detailed analysis of the predicted structure (no Pam2CSK4 docked) shows that the cavity is not located within the LRR solenoid but is created by a long loop wrapped around the main body of the LRR domain (See Figure 5 in Discussion). Thus, despite a similar volume of cavity and overlapping location, this may be an example of convergent evolution of cavities in iTLRs.

*Ciona intestinalis* is closer to vertebrates in the phylogenetic tree; thus, ciTLR2 could be the iTLR closest to the vTLR lineage. In contrast to LeTLR2, ciTLR2 has a typical iTLR signature with an additional LRR domain separated by a characteristic C-terminal LRR repeat. In an unusual feature, its N-terminal LRR is broken into two parts with a break in the hydrogen bond network of the concaved beta sheet. It also contains a cavity with a volume of 1566 Å^3^ that is within the vTLR range and has a paralog with a smaller cavity measuring 748 Å^3^. However, similar to LeTLR2, the cavity is not inside the LRR solenoid but is an artifact of the break between the two halves of the N-terminal LRR domain (Figure 2B, Figure 5). It is possible to argue that this is an ancestral form of the vTLR cavity; however, given its broad specificity and unclear phylogenetic position [33], it is more likely that it is not a bona fide ortholog of the vTLRs.

We also investigated the presence of potential TLR2 orthologs in other invertebrates in the phylum Chordata, as they are the closest invertebrate relatives to vertebrates. Among all TLRs identified in *Branchiostoma, Saccoglossus*, and *Strongylocentrotus*, none had sizable cavities in their LRR domains.

As discussed earlier, several species (within the manually analyzed group) have TLR2 paralogs. The cavities were similar in shape and position along the chain, indicating their origin of duplication (see Table 1 in supp. info). The clawed frog TLR2 was further analyzed by docking with PAM2CSK4 to illustrate that duplication results in the functional conservation of ligand binding. Paralogs were also docked to LTA, and the results are shown in Table 6 of Supp. Info. In vTLRs species duplicates are relatively common; approximately a dozen species (According to the Ensembl Gene gain/loss tree TLR2) had at least one TLR2 paralog, suggesting continuous ancestral evolution adapting to different environments and pathogens via gene duplication.

### Comparison of cavities of all 239 vTLR AF3 models (Tables 2, 3, and 4 in Supp. Info)

The analysis of 239 vTLR cavity volumes was conducted using two different tools, PyVOL (Table 10) and Fpocket 4 (Table 11), to demonstrate the consistency of the volume calculation. Fpocket4 also calculates other cavity parameters. The results show a comparison of specific parameters, including volume, in five commonly used groups of animals: Fish, Amphibians, Reptiles, Birds and Mammals (Tables 2, 3 and 4 provided in Supp. Info).

Table 2 shows that the average volume and Standard Deviation (SD) of vTLRs, as calculated by PyVOL and Fpocket4, were 1197 ± 45.66 and 1231 ± 49.63 Å^3^, respectively. This indicates that the vTLR volume calculations were consistent and measurable using both tools. Table 2 also shows the average volume and SD calculated by Fpocket4 and PyVOL for each of the five classes (fish, amphibians, reptiles, birds and mammals). Except for amphibians, all class volumes calculated by PyVOL and Fpocket4 were within the SD of the average measurements for all 239 vTLRs genes. This discrepancy for amphibians arises because only two of the 239 human vTLR orthologs, which were two Xenopus duplicates, were annotated in the Ensembl Database. These volumes were 1455.48 (LDO) and 1705.16 (paralogs) Å^3^, compared to the average of 1197 ± 45.66 Å^3^ for the vTLRs.

Table 3 shows the average and SD of the 20 parameters calculated using Fpocket 4 for all five groups. Table 4 shows that the TLR2 cavity volume and most of the other features did not differ significantly among vertebrate classes (F = 1.20, P = 0.32) according to one-way ANOVA, indicating a high degree of consistency in the overall pocket size across groups. This further supports the ancestral consistency fundamental to the size, shape, polarity, and ligand-accommodating capacity of the vTLR pocket in all five animal groups.

Among the 20 parameters analyzed, flexibility emerged as the most significant differentiator (p < 0.001), identifying fish as the distinct outlier with the most rigid pockets (mean 0.638) compared to the significantly more flexible pockets of birds (0.751) and reptiles (0.743), whereas mammals (0.689) retained rigidity closer to that of fish. Alpha Sphere Density (p =0.02) revealed further divergence, specifically contrasting birds, which exhibited the highest density (8.88), against reptiles, which showed the lowest (7.70). Additionally, the Center of Mass Distance (p = 0.03) indicated significant variance in pocket shape, where reptiles possessed the most compact, centered pockets (17.70) in contrast to birds, which displayed the most elongated or distributed binding sites (22.50)

These findings suggest that vertebrate evolution has preserved the pocket size and physicochemical properties while selectively tuning the pocket flexibility and internal spatial arrangement, likely reflecting adaptation to the distinct pathogen repertoires encountered by different lineages in their respective environments. Amphibian values showed greater variance for some parameters, although inferences for this class were limited by small sample size. Overall, the results support a model in which the vTLR pocket volume and chemistry are evolutionarily conserved, whereas the internal geometry and flexibility are modulated in a lineage-specific manner appropriate for their respective environmental pathogen repertoires.

In addition to comparing the cavity structures, virtual ligand docking was performed to further assess cavity function and structure (Figure 4). The primary function of the large cavity in TLR2 is to bind various ligands from gram-positive bacteria, mycobacteria, mycoplasmas and fungi. Pam2CSK4 binding (Figure 4) illustrates that the function of the cavity is consistent among vertebrates, reinforcing the notion that this cavity is an ancestral feature. The binding consistency in invertebrates, such as LeTLR2, reinforces the retention of the convergent function of this binding cavity. Species TLR2 duplications (Figure 4) were consistent across all species. The binding affinity was calculated to further support this finding for all species and their respective duplicates. The binding affinities were remarkably similar, with -13.7582 ± 0.408, all within the same standard deviation. Thus, it can be demonstrated that ligand binding reinforces the consistency of the structure and binding of the common and significant gram-positive ligand Pam2CSK4.

In Figure 4 **(A)**, the clawed frog TLR2 (LDO) and **(B)** its paralog are shown bound to Pam2CSK4, with only the binding cavity highlighted in green. The binding energies of the clawed frog and its TLR2 paralog gene bound to Pam2CSK4 were -14.02 and 13.90 kcal/mol, respectively. These values were nearly identical, with an average of -13.7582 ±0.408 kcal/mol. Ligand binding and similar free energy binding values illustrate how, within vertebrate species duplicates, the ligand-binding function of the cavity is consistent and conserved, reinforcing the evidence of ancestral (divergent) evolution. Another comparison shown **(C)** Human TLR2 and **(D)** Lamprey TLR2 binding to the TLR2 agonist Pam2CSK4, with their respective central cavities highlighted in green. The binding energies of human and Lamprey TLR2 to Pam2CSK4 were -13.77 and -13.94 kcal/mol, respectively. These values are similar, and the average is -13.7582 ±0.408 kcal/mol. Ligand binding and similar free energy binding values illustrate how divergent vertebrate species, such as lampreys and humans, conserve the ligand-binding function of the cavity, reinforcing the evidence of the ancestral (divergent) evolution of this structure. The consistency of ligand binding between humans and lampreys further reinforces Lamprey TLR2 as the ancestor of all vTLR via divergent evolution. Another comparison **(E)** LeTLR2 is shown to be bound to Pam2CSK4, with their respective central cavities highlighted in green.

The binding energies of human and LeTLR2 to Pam2CSK4 were -13.77 and -14.06 kcal/mol, respectively. The presence of a large cavity, which conserves ligand-binding function to TLR2, supports that LeTLR2 could be a TLR2 ortholog; however, Pam2CSK4 binds differently than when bound to vTLRs with its long fatty acid palmitoyl chains oriented in opposite directions, which may be an artifact due to Boltz 1 (it also appeared in Boltz 2[34] preprint). The cavity embedded in the solenoid structure only appeared when the Pam2CSK4 ligand was bound to LeTLR2 (Figure 2A). In addition, the misalignment and off-centered cavity seem to reinforce that LeTLR2 is not a functional ortholog but a homolog of vTLR.

Ligand binding was used to demonstrate that the convergent TLR2 cavity observed in leeches retained its functional capacity to bind LTC and Pam2CSK4. Both ligands were bound to LeTLR2, and their binding patterns were similar to those observed in vertebrate complexes. Furthermore, their binding was within the standard deviations of both vertebrate groups. Ligand binding to LTA is shown in the Supp. Info (Table 6).

TLR2, TLR1, and TLR6 primarily exist as monomers in the plasma membrane. A generally accepted mechanism of TLR2 is that a ligand docks to the monomeric TLR2 and stabilizes the appropriate heterodimer (TLR1/2 or TLR2/6). The docking, as described above and shown in Figure 4, captures the first phase of the cooperative nature of the ligand-binding requirements. Table 7 in Supp. Info show that the modeled AF3 and PDB of Human and Mouse dimers of TLR2/1/6 docked with diacyl and triacyl lipopeptides (Pam2/3CSK4) showed little difference in the volume of the cavity; thus, dimerization and docking should not change the volume substantially, allowing ligand docking to be investigated as an illustrative exploration and good model. In addition, the similarity of the cavity volumes in the AF3 and PDB models shows that recent AI protein folding tools are excellent for volume cavity modeling and allow consistent measurement analyses.

Additionally, it is crucial to recognize that the palmitoyl fatty acid chains of LTC and Pam2CSK4 directly interact with the large cavity in TLR2, rather than with the head groups. Tri-acylated lipopeptides interact with the TLR2/TLR1 complex as the third fatty acid palmitoyl group docks into the cavity of TLR1. TLR2 has the largest cavity of the TLR family, although TLR1/7/8/9 have cavity features that are much smaller and bind different agonists. To accurately dock ligands, a heterodimer with TLR6 or TLR1 must be employed, necessitating further investigation into the orthologs of TLR6 and TLR1 across vertebrate species and their respective duplicates. This is a potential direction for future research.

Although the CiTLR2 gene may represent the earliest emergence of a TLR2-like form and the earliest iTLR, the presence of an additional small LRR domain, which is absent in all vTLRs, indicates that the invertebrate ancestor of all vTLRs must have undergone significant changes, such as domain deletion, resulting in the loss of the secondary LRR domain. Further investigation into the splicing of this specific CiTLR2 gene may provide additional clarification. The CiTLR2 secondary LRR domain spans five exons; thus, a single exon deletion is not possible, and multiple exon deletions are possible but less likely to occur. Thus, CiTLR2 is likely a TLR homolog with a cavity that emerged via convergent evolution. However, while more data and *Ciona* genomes become available, at this point lampreys and hagfish remain the primary point of comparison for vertebrate evolution of TLR2 lineage.

## Discussion

Analysis of vTLR2 cavity parameters across all vertebrates provides strong evidence for the ancestral origin of this cavity at the base of the vertebrates. The consistency in key parameters calculated by Fpocket4 (Tables 2–4), such as volume, hydrophobicity, polarity, and ligand-accommodating capacity, across fish, amphibians, reptiles, birds, and mammals underlines the conservation of core structural and chemical features since the earliest vertebrate ancestors, such as lampreys. The lack of significant differences in most pocket parameters, despite the vast evolutionary distances between these groups, suggests that the basic architecture and function of the TLR2 cavity were established early and have been maintained because of their essential roles in innate immune recognition and ligand-binding. Simultaneously, observed lineage-specific variations in pocket flexibility, packing, and internal geometry point to adaptive fine-tuning, allowing each group to optimize its immune function against unique pathogen landscapes. In addition, fine-tuning to the respective environment may also be observed in gene duplication in approximately a dozen vertebrate genera (Figure 3). Together, these findings support a model in which vertebrate TLR2s have preserved a common ancestral pocket design, whereas evolutionary pressures have selectively shaped their dynamic properties to address specific environmental challenges. Investigating the origins of innate immune recognition mechanisms, such as the recognition of gram-positive bacterial ligands by TLR2, which are critical in the host response to infectious diseases, may enhance our understanding of immune system evolution and identify conserved structural features underlying pathogen recognition across metazoans (Figure 4).

**Figure 3:**
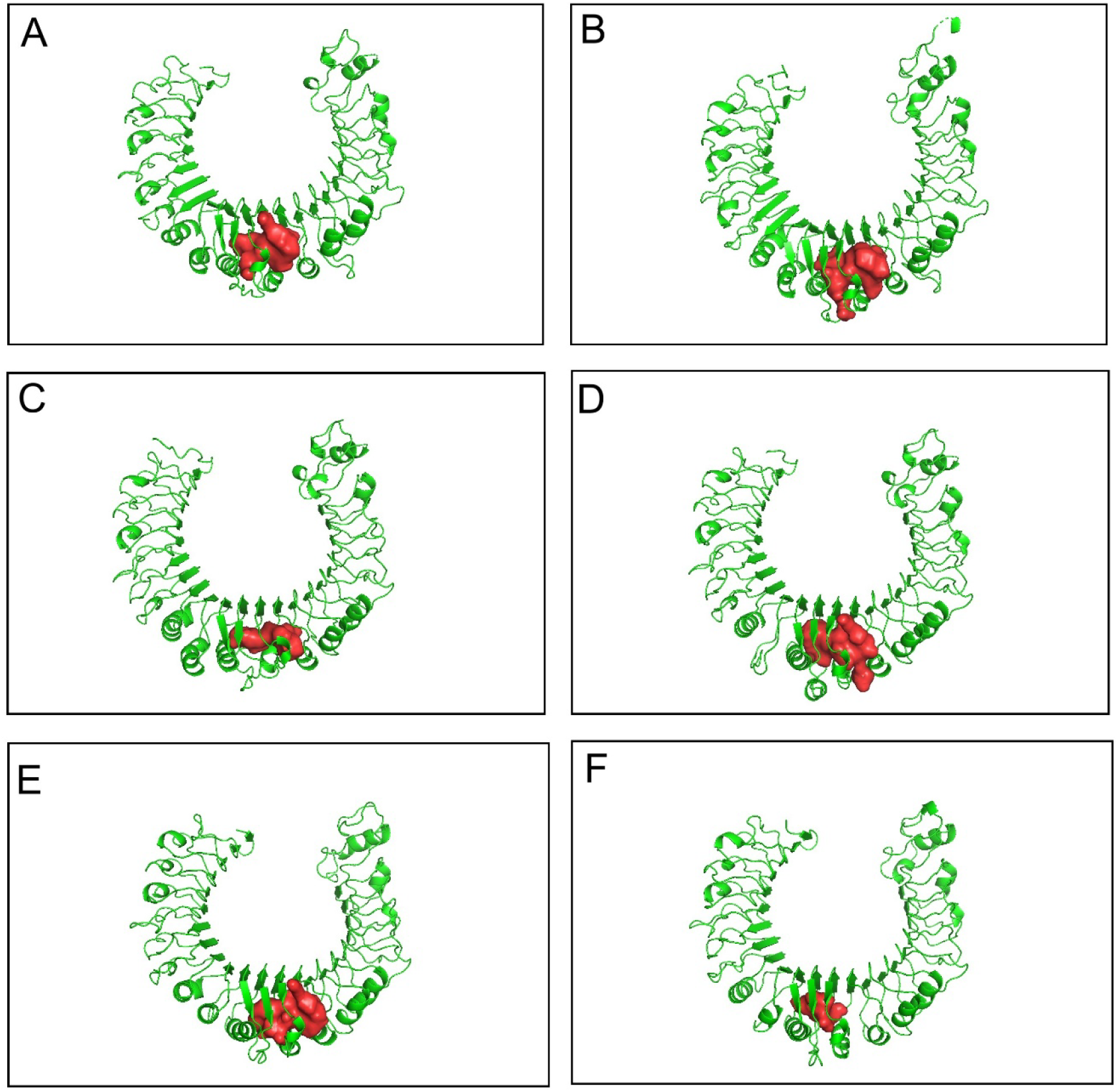
shows a comparison of several TLR2 in species paralogs: clawed frog (*Xenopus laevis*) **(A)** LDO, **(B)** duplicate, chicken (*Gallus gallus domesticus*) **(C)** LDO (**D)** duplicate, and green anole (*Anolis carolinensis*) **(E)** LDO **(F)** duplicate.

**Figure 4:**
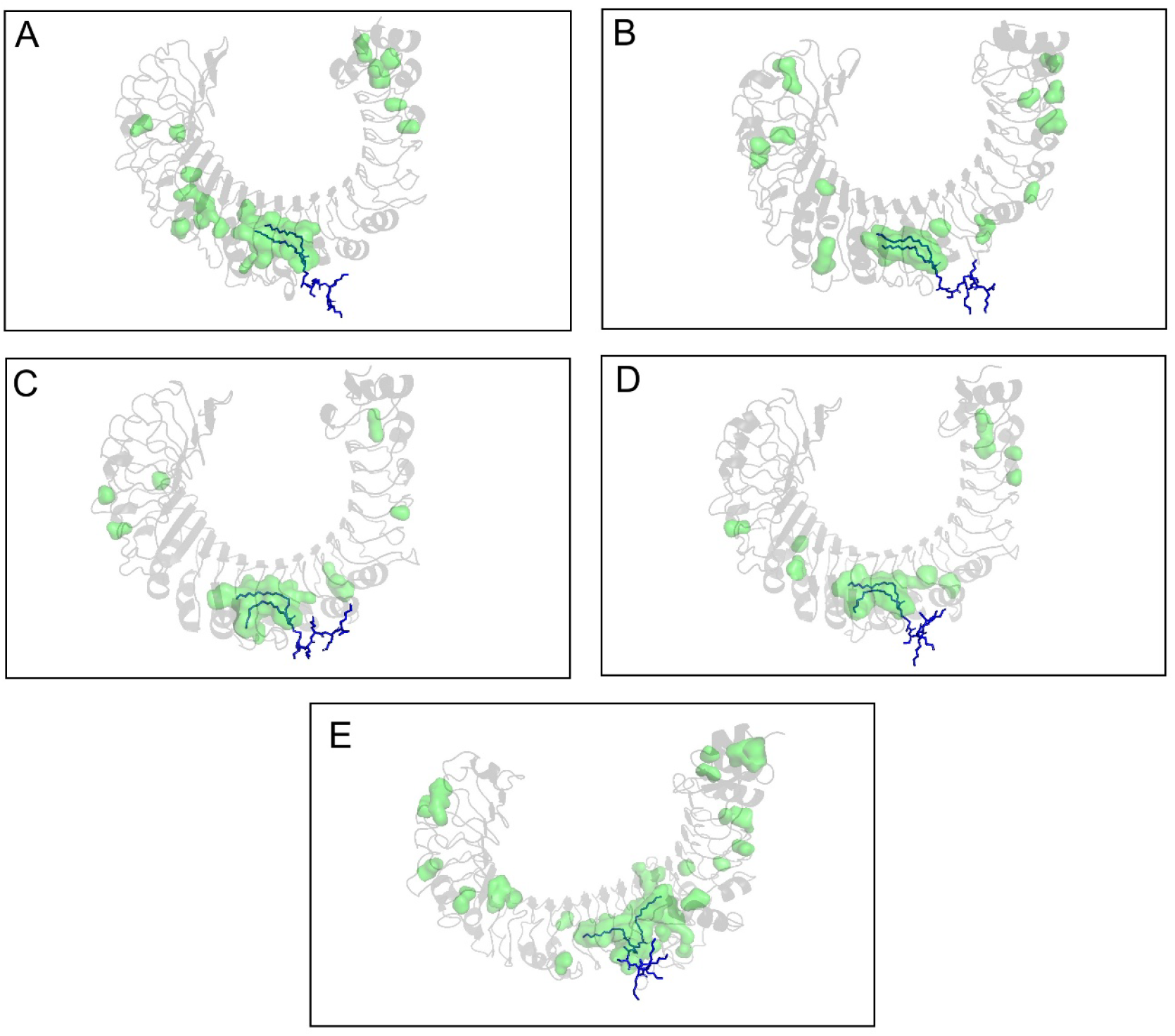
Ligand docking of diacyl lipopeptide synthetic PAM2CSK4 across the vTLRs. (Table 5 in Supp. Info) **(A)** human **(B)** lamprey, **(C)** clawed frog (LDO), (D) clawed frog (duplicate) and **(E)** leech.

**Figure 5:**
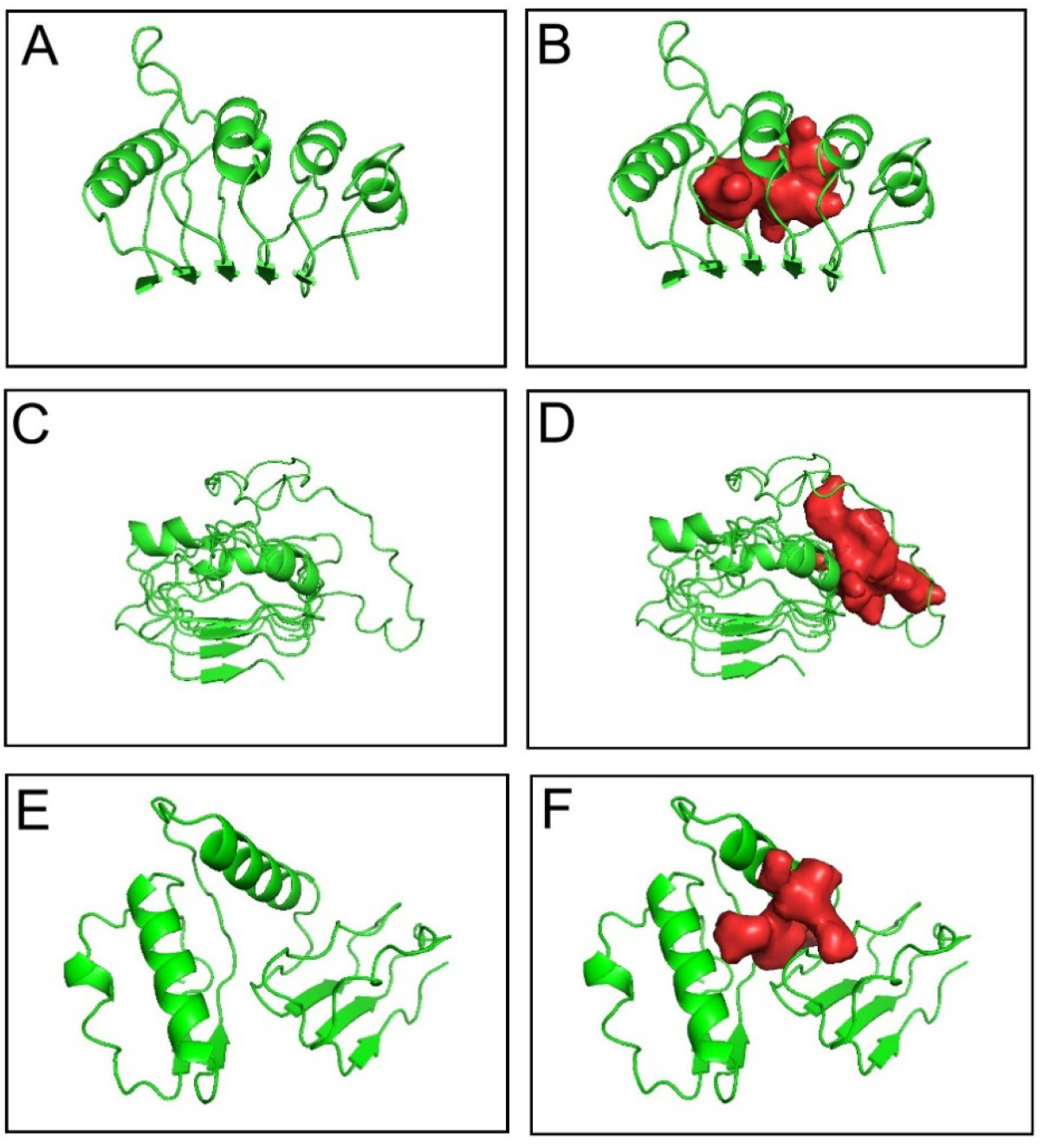
Structural comparison of cavity carrying section of the LRR domain in a sample vertebrate TLR2 (human) and two invertebrate putative TLR2s. **(A)** Human TLR2 segment shown with no cavity, and **(B)** the same segment with the cavity shown to be embedded deep in LRR solenoid. LeTLR2 **(C)** segment of the LRR domain showing a large loop adjacent to solenoid and **(D)** the cavity as formed between loop and the main solenoid body. **(E)** CiTLR2 has a break between two segments of the N-terminal LRR domains with broken H-bond pattern, **(F)** CiTLR2 cavity is formed between breaks in LRR domains.

Invertebrate LeTLR2 and CiTLR2 cavities have volumes well within the distribution of vTLRs; however, the exact position of the cavity in relation to the LRR solenoid – outside of the solenoid, formed by a large loop or a break within the domain. Together with the other invertebrate-specific features of the entire protein, this suggests that it emerged through a process of convergent evolution or perhaps represents an early form of the cavity. The relative dearth of genomic data on deuterostome invertebrates does not allow a deeper analysis of how common this cavity is in this group. Lamprey and hagfish TLR2s may be the most direct descendants of the original ancestral vTLR cavity in vertebrates.

We relied on an existing phylogenetic tree from PANTHERDB and ortholog annotations from UniProt and Ensembl databases. PANTHERDB and Ensembl contain only representative species from some genera; therefore, there is only one representative of lampreys (*Petromyzon marinus*) and hagfish (*Eptatretus burgeri*). Given the importance of these species in the early evolution of the TLR2 cavity, we extended our analysis to other hagfish and lamprey species. Analysis of TLR2s from Atlantic hagfish (HgTLR2) (*Myxine glutinosa*) and inshore hagfish (*Eptatretus burgeri*) revealed that they do, in fact, contain a vTLR-like cavity and can dock Pam2CSK4. Several other full genomes of lampreys exist in NCBI and were investigated, it was confirmed that all complete, high coverage genomes of other lamprey species contained TLR2s orthologues.

Therefore, the most likely scenario is that hagfish (Myxini) progressively lost the TLR2 cavity feature over time, as some hagfish species (Atlantic and inshore) still contain it. (S2 in Supp. Figures showing Pam2CSK4 docking into Inshore and Atlantic hagfish are shown in the Supp. Info).

In addition, all LRR-containing proteins (n = 325) in the human proteome were analyzed using both Fpocket4 and PyVOL (the tables of values are provided in Table 8 & 9 respectively). While there were no cavities matching those in TLR2, two were worth noticing: Negative regulator of reactive oxygen species (NRROS) and NACHT, LRR, and PYD domains-containing protein 3 (NLRP3). The cavity in the latter has been discussed in the literature [12], whereas the structure of the former has never been experimentally analyzed.

LeTLR2 also appeared to be in a similar 300–450 residue position near the middle of the LRR domain. However, the cavity is not formed by the repeats a located inside the solenoid, as it is in vTLR2s, domain but mostly by a unusually long loop; LeTLR2 also failed Fpocket4 to generate a cavity embedded in the solenoid structure without the Pam2CSK4 ligand docked. Despite the similar volume of the cavity generated by binding, this could be an artifact of ligand binding using Boltz 1 and 2 modeling. It has been previously reported that an alternative theory to convergent evolution is gene transfer, as leeches feed on blood (and white blood cells expressing TLR genes) and gain vTLR by parasitizing vertebrates [35]. Based on these analyses we believe that LeTLR2, as a member of the TLR family is a TLR2 homolog but is not a TLR2 ortholog.

Finally, CiTLR2 did not align well with LeTLR2, HgTLR2, human, or lamprey TLR2 (Figure 3C Supp. Figures). CiTLR2 was unable to produce a Pam2CSK4 Model in Boltz 1 or 2; thus, while it contains a cavity in its LRR domain, whether the classical di-acyl ligand binding feature is conserved is not known, although several experimental studies have shown that TLR2 agonists upregulate TLR2 genes in *Ciona* sp. However, this does not indicate the physical docking of ligands to CiTLR2. As previously mentioned, the cavity found in CiTLR2 is positioned at the breakpoint of two LRR domains in the protein, in contrast to the centrally embedded cavity in the solenoid of vTLRs. Thus, the structure, cavity location, and alignment mismatch all suggest that CiTLR2 may be an example of convergent evolution and the earliest appearance of a cavity in a TLR family and is not a TLR2 ortholog.

Structural phylogeny can thus be employed to investigate consistent features appearing in invertebrates and vertebrates, such as the cavity feature in vTLR, to determine origin, evolutionary trends, and function. The existence of true vTLR orthologs in invertebrates remains uncertain as the annotated “TLR” genes (iTLR) in the families *Ciona, Helobdella, Branchiostoma, Saccoglossus*, and *Strongylocentrotus*, none were found. Homologs of TLR2 exist in the iTLR group as evident by LeTLR2 and CiTLR2. Currently, the unique TLR2 cavity feature present in vTLR most likely originated in Agnatha, and specifically in a common ancestor of the current extant Hagfish and Lamprey species. The TLR2 cavity is consistent in shape and function across vTLRs, although minor variations exist as a means of adaptation to local environmental pathogens. The presence of this cavity throughout vertebrates is the result of divergent (ancestral) evolution that extends from the base of an ancient member of Agnatha.

## Acronyms

PAMPs: pathogen associated molecular patterns
DAMPs: danger associated molecular patterns
TIR: Toll/interleukin-1 receptor domain
LDO: Least Diverged Ortholog
CiTLR2: *Ciona intestinalis* putative TLR2 (F6SGF2_CIOIN)
LeTLR2: *Helobdella robusta* putative TLR2 (T1EUA2_HELRO)
HgTLR2: *Myxine glutinosa* (Atlantic hagfish) TLR2 (XP_067991900.1)
vTLR: (vertebrate TLRs)
iTLR: (invertebrate TLRs)

## Tables attached as separate documents

**Table 1.** Curated vertebrate TLR2 genes and their species paralogs (duplications)

**Table 2:** vTLR Pocket Volume Summary

**Table 3.** FPocket Parameter Statistics by Vertebrate Group

**Table 4.** One-way ANOVA of fpocket Parameters Across Vertebrates

**Table 5:** Pam2CSK4 (Di-acylated Lipo-peptide) Binding Analysis

**Table 6:** LTA Binding Affinity Analysis

**Table 7:** TLR2 Cavity parameters in complex with/out TLR6 or TLR1 and/or Pam2/3CSK4 (Di-acyl or Tri-acyl) lipopeptide

**Table 8:** All 22 Fpocket4 parameters calculated from each 326 Human proteome LRR (“Leucine Rich Repeat”) proteins. Fpocket 4 provides 22 descriptors that characterize pockets based on their geometry, hydrophobicity, and chemical environment. Key physical indicators include Volume, and others, which together define the cavity’s shape and suitability for ligand interaction.

**Table 9:** PyVOL volume calculations in A^3^ for each 326 Human LLR proteins

**Table 10:** PyVOL volume calculations in A^3^ for each 239 vTLR orthologs from Ensembl

**Table 11:** All 22 Fpocket4 parameters calculated from each 239 vTLR orthologs from Ensembl. These parameters are the same as Table 8. Fpocket 4 provides 22 descriptors that characterize pockets based on their geometry, hydrophobicity, and chemical environment. Key physical indicators include Volume, and others, which together define the cavity’s shape and suitability for ligand interaction.

## References

1. Di Lorenzo A, Bolli E, Tarone L, Cavallo F, Conti L. Toll-Like Receptor 2 at the Crossroad between Cancer Cells, the Immune System, and the Microbiota. IJMS. 2020;21: 9418. doi:10.3390/ijms21249418

2. Kang JY, Nan X, Jin MS, Youn S-J, Ryu YH, Mah S, et al. Recognition of Lipopeptide Patterns by Toll-like Receptor 2-Toll-like Receptor 6 Heterodimer. Immunity. 2009;31: 873– 884. doi:10.1016/j.immuni.2009.09.018

3. Lee JO, Jin MS, Kim SE, Heo JY. Crystal structure of the TLR1-TLR2 heterodimer induced by binding of a tri-acylated lipopeptide. 2007. doi:10.2210/pdb2z80/pdb

4. Oliveira-Nascimento L, Massari P, Wetzler LM. The Role of TLR2 in Infection and Immunity. Front Immun. 2012;3. doi:10.3389/fimmu.2012.00079

5. Jumper J, Evans R, Pritzel A, Green T, Figurnov M, Ronneberger O, et al. Highly accurate protein structure prediction with AlphaFold. Nature. 2021;596: 583–589. doi:10.1038/s41586-021-03819-2

6. Abramson J, Adler J, Dunger J, Evans R, Green T, Pritzel A, et al. Accurate structure prediction of biomolecular interactions with AlphaFold 3. Nature. 2024;630: 493–500. doi:10.1038/s41586-024-07487-w

7. Krishna R, Wang J, Ahern W, Sturmfels P, Venkatesh P, Kalvet I, et al. Generalized biomolecular modeling and design with RoseTTAFold All-Atom. Science. 2024;384. doi:10.1126/science.adl2528

8. Lin Z, Akin H, Rao R, Hie B, Zhu Z, Lu W, et al. Evolutionary-scale prediction of atomic-level protein structure with a language model. Science. 2023;379: 1123–1130. doi:10.1126/science.ade2574

9. Hayes T, Rao R, Akin H, Sofroniew NJ, Oktay D, Lin Z, et al. Simulating 500 million years of evolution with a language model. Science. 2025;387: 850–858. doi:10.1126/science.ads0018

10. Lokhande KB, Singh A, Vyas R, Joe S, Asthana S, Pawar K. 5′-tRNAHisGUG fragment: A preferred endogenous TLR7 ligand with reverse sequence activation insights. Biophysical Journal. 2025;124: 1961–1978. doi:10.1016/j.bpj.2025.04.027

11. Beesu M, Caruso G, Salyer ACD, Khetani KK, Sil D, Weerasinghe M, et al. Structure-BasedDesign of Human TLR8-Specific Agonists with Augmented Potency andAdjuvanticity. J Med Chem. 2015;58: 7833–7849. doi:10.1021/acs.jmedchem.5b01087

12. Ma Q. Pharmacological Inhibition of the NLRP3 Inflammasome: Structure, Molecular Activation, and Inhibitor-NLRP3 Interaction. Pharmacological Reviews. 2023;75: 487–520. doi:10.1124/pharmrev.122.000629

13. Tang J, Han Z, Sun Y, Zhang H, Gong X, Chai J. Structural basis for recognition of an endogenous peptide by the plant receptor kinase PEPR1. Cell Res. 2014;25: 110–120. doi:10.1038/cr.2014.161

14. Pan J, Ahmad MZ, Zhu S, Chen W, Yao J, Li Y, et al. Identification, Classification and Characterization Analysis of FBXL Gene in Cotton. Genes. 2022;13: 2194. doi:10.3390/genes13122194

15. Newman RM, Salunkhe P, Godzik A, Reed JC. Identification and Characterization of a Novel Bacterial Virulence Factor That Shares Homology with Mammalian Toll/Interleukin-1 Receptor Family Proteins. Infect Immun. 2005;74: 594–601. doi:10.1128/iai.74.1.594-601.2006

16. Zhang Y, Zagnitko O, Rodionova I, Osterman A, Godzik A. The FGGY Carbohydrate Kinase Family: Insights into the Evolution of Functional Specificities. PLoS Comput Biol. 2011;7: e1002318. doi:10.1371/journal.pcbi.1002318

17. Adiba S, Nizak C, Van Baalen M, Denamur E, Depaulis F. From grazing resistance to pathogenesis: the coincidental evolution of virulence factors. PLoS ONE. 2010;5: e11882. doi:10.1371/journal.pone.0011882

18. Sacristán S, Goss EM, Eves-Van Den Akker S. How Do Pathogens Evolve Novel Virulence Activities? MPMI. 2021;34: 576–586. doi:10.1094/mpmi-09-20-0258-ia

19. Dyer SC, Austine-Orimoloye O, Azov AG, Barba M, Barnes I, Barrera-Enriquez VP, et al. Ensembl 2025. Nucleic Acids Research. 2024;53: D948–D957. doi:10.1093/nar/gkae1071

20. Bateman A, Martin M-J, Orchard S, Magrane M, Adesina A, Ahmad S, et al. UniProt: the Universal Protein Knowledgebase in 2025. Nucleic acids research. 2024;53: D609–D617. doi:10.1093/nar/gkae1010

21. Wohlwend J, Corso G, Passaro S, Getz N, Reveiz M, Leidal K, et al. Boltz-1 Democratizing Biomolecular Interaction Modeling. bioRxiv. 2025;28. doi:10.1101/2024.11.19.624167

22. Neurosnap—Computational Biology, Simplified [Internet]. [cited 26 Jan 2026]. Available: https://neurosnap.ai/.

23. Vangone A, Schaarschmidt J, Koukos P, Geng C, Citro N, Trellet ME, et al. Large-scale prediction of binding affinity in protein-small ligand complexes: the PRODIGY-LIG web server. Bioinformatics. 2018;35: 1585–1587. doi:10.1093/bioinformatics/bty816

24. Edgar RC. MUSCLE: multiple sequence alignment with high accuracy and high throughput. Nucleic Acids Research. 2004;32: 1792–1797. doi:10.1093/nar/gkh340

25. Smith RHB, Dar AC, Schlessinger A. PyVOL: a PyMOL plugin for visualization, comparison, and volume calculation of drug-binding sites. Cold Spring Harbor Laboratory; 2019. doi:10.1101/816702

26. Schmidtke P, Le Guilloux V, Maupetit J, Tufféry P. fpocket: online tools for protein ensemble pocket detection and tracking. Nucleic Acids Research. 2010;38: W582–W589. doi:10.1093/nar/gkq383

27. Kochnev Y, Durrant JD. FPocketWeb: protein pocket hunting in a web browser. J Cheminform. 2022;14. doi:10.1186/s13321-022-00637-0

28. Takkouche A, Ichii K, Qiu X, Jaroszewski L, Godzik A. Divergence of the Individual repeats in the leucine-rich repeat domains of human Toll-like receptors explain their diversity and functional adaptations. Cold Spring Harbor Laboratory; 2024. doi:10.1101/2024.09.30.615863 (Preprint)

29. Liu G, Zhang H, Zhao C, Zhang H. Evolutionary History of the Toll-Like Receptor Gene Family across Vertebrates. Genome Biology and Evolution. 2019;12: 3615–3634. doi:10.1093/gbe/evz266

30. Pergolizzi S, Fumia A, D’Angelo R, Mangano A, Lombardo GP, Giliberti A, et al. Expression and function of toll-like receptor 2 in vertebrate. Acta Histochemica. 2023;125: 152028. doi:10.1016/j.acthis.2023.152028

31. Thomas PD. GIGA: a simple, efficient algorithm for gene tree inference in the genomic age. BMC Bioinformatics. 2010;11. doi:10.1186/1471-2105-11-312

32. Thomas PD, Ebert D, Muruganujan A, Mushayahama T, Albou L, Mi H. PANTHER: Making genome-scale phylogenetics accessible to all. Protein Science. 2021;31: 8–22. doi:10.1002/pro.4218

33. Sasaki N, Ogasawara M, Sekiguchi T, Kusumoto S, Satake H. Toll-like Receptors of the Ascidian Ciona intestinalis: PROTOTYPES WITH HYBRID FUNCTIONALITIES OF VERTEBRATE TOLL-LIKE RECEPTORS. Journal of Biological Chemistry. 2009;284: 27336– 27343. doi:10.1074/jbc.m109.032433

34. Passaro S, Corso G, Wohlwend J, Reveiz M, Thaler S, Somnath VR, et al. Boltz-2: Towards Accurate and Efficient Binding Affinity Prediction. bioRxiv. 2025;53. doi:10.1101/2025.06.14.659707

35. Silver A, Graf J. Innate and procured immunity inside the digestive tract of the medicinal leech. Invertebrate survival journal : ISJ. 2011;8.

